# DOCK3 regulates normal skeletal muscle regeneration and glucose metabolism

**DOI:** 10.1101/2023.02.22.529576

**Authors:** Adrienne Samani, Muthukumar Karuppasamy, Katherine G. English, Colin A. Siler, Yimin Wang, Jeffrey J. Widrick, Matthew S. Alexander

## Abstract

DOCK (dedicator of cytokinesis) is an 11-member family of typical guanine nucleotide exchange factors (GEFs) expressed in the brain, spinal cord, and skeletal muscle. Several DOCK proteins have been implicated in maintaining several myogenic processes such as fusion. We previously identified DOCK3 as being strongly upregulated in Duchenne muscular dystrophy (DMD), specifically in the skeletal muscles of DMD patients and dystrophic mice. *Dock3* ubiquitous KO mice on the dystrophin-deficient background exacerbated skeletal muscle and cardiac phenotypes. We generated *Dock3* conditional skeletal muscle knockout mice (*Dock3* mKO) to characterize the role of DOCK3 protein exclusively in the adult muscle lineage. *Dock3* mKO mice presented with significant hyperglycemia and increased fat mass, indicating a metabolic role in the maintenance of skeletal muscle health. *Dock3* mKO mice had impaired muscle architecture, reduced locomotor activity, impaired myofiber regeneration, and metabolic dysfunction. We identified a novel DOCK3 interaction with SORBS1 through the C-terminal domain of DOCK3 that may account for its metabolic dysregulation. Together, these findings demonstrate an essential role for DOCK3 in skeletal muscle independent of DOCK3 function in neuronal lineages.

## Introduction

Skeletal muscle is essential for the body’s locomotive function, maintenance of the skeleton structure, and it retains a trademark capacity for repair and regeneration(Chaillou and Lanner 2016). As an organ, skeletal muscle plays a major role in the processing and utilization of glucose in response to insulin. Through this mechanism, it is responsible for approximately 80% of postprandial glucose uptake from circulation, making it critical to maintaining metabolic homeostasis at the organismal level(DeFronzo and Tripathy 2009). Many key cell signaling pathways are essential for normal muscle cell regeneration, migration, membrane fusion, repair, and muscle metabolism during growth and development(Sampath, Sampath et al. 2018). Several Rho GTPases function as molecular switches during cell signaling pathways important to the regulation of the F-actin cytoskeleton(Noviello, Kobon et al. 2021). Additional downstream Rho signaling effectors, such as RAC1 and CDC42, have been implicated in myogenic processes including myogenic differentiation, fusion, myoblast proliferation, and are known to influence the regenerative capacity within the skeletal muscle(Samson, Will et al. 2007).

The *DOCK* gene family is an 11-member class of guanine nucleotide exchange factors capable of influencing multiple pathways involved in cellular fusion, migration, and survival in a myriad of tissue types(Côté and Vuori 2002). Many of these DOCK proteins are highly expressed in the brain, spinal cord, and muscle(Aguet, Anand et al. 2020). Recent studies have demonstrated that DOCK proteins play essential functional roles in important skeletal muscle processes in health and disease(Samani, English et al. 2022). For example, DOCK1 and DOCK5 have been illustrated as crucial players in myoblast fusion(Moore, Parkin et al. 2007, Laurin, Fradet et al. 2008). Along those same lines, DOCK3 plays a key role in RAC1 activation and WAVE signaling in neurons and skeletal muscle(Namekata, Enokido et al. 2004, Namekata, Harada et al. 2010, Helbig, Mroske et al. 2017). Patients with loss-of-function *DOCK3* variants present with a variety of developmental disorders such as intellectual disability, developmental delay, ataxia, and muscle hypotonia(Helbig, Mroske et al. 2017, Iwata-Otsubo, Ritter et al. 2018).

Previously, we identified DOCK3 as a dosage-sensitive biomarker of DMD in which disease severity correlated with increased DOCK3 expression in the skeletal muscles of affected patients and dystrophic *mdx*^*5cv*^ mice(Reid, Wang et al. 2020). Interestingly, the adult skeletal muscle of the ubiquitous *Dock3* KO mice showed a reduction in myofiber diameter and overall structure, reduced muscle mass, and metabolic dysfunction. DOCK3 is expressed in both the central nervous system and in skeletal muscle, thus we sought to understand the role of DOCK3 exclusively in the skeletal muscle by generating a skeletal muscle-specific conditional mouse knockout of *Dock3* within the myofiber. We hypothesized that a muscle-specific loss of DOCK3 would disrupt major myogenic processes and protein-protein interactions, subsequently undermining muscle regeneration, metabolism, and overall muscle function. We generated a muscle-specific mouse model (henceforth referred to as *Dock3* mKO) to understand the role of DOCK3 in overall muscle health. We evaluated *Dock3* mKO mouse models and found mild disruptions in muscle integrity and function using activity tracking, but no evidence of contractile deficits using *ex vivo* functional assays. We evaluated the role of DOCK3 in muscle repair and showed an impairment in the skeletal muscle’s capacity to repair in the *Dock3* mKO mice following a cardiotoxin-induced injury. Finally, we evaluated the impact of DOCK3 on glucose metabolism via its activation of the GLUT4 transporter and identified a novel protein-protein interaction with the insulin adaptor protein Sorbin and SH3 domain containing 1 (SORBS1). We demonstrated that DOCK3 is essential for normal skeletal muscle regeneration and metabolic regulation within the skeletal muscle.

## Materials and Methods

### Animals

*Dock3* conditional knockout mice were generated commercially (Cyagen; Santa Clara, CA). These mice were generated using a CRISPR-Cas9 approach to generate an out-of-frame *Dock3* deficient mouse upon *Cre*-mediated recombination by excision of exons 8 and 9 of the mouse *Dock3* transcript (NCBI Reference Sequence: NM_153413). Guide RNAs (gRNAs) targeting the intronic regions flanking mouse *Dock3* exons 8 and 9 were used along with a homologous recombination vector were injected into wild type *C57BL/6* mouse embryos (Taconic Biosciences; Germantown, NY) to generate Dock3 conditional knockout mice (*Dock3*^*fl/fl*^). The targeting homologous recombination vector contained loxP sites flanking mouse *Dock3* exons 8 and 9 and was co-injected with the gRNAs and Cas9 mRNA. F_0_ founder mice were identified by PCR followed by sequence analysis and were then backcrossed to wild type mice to test germline transmission and F_1_ animal generation. PCR oligonucleotide primers used to genotype the genomic tail DNA from isolated biopsies from the loxP sites in the *Dock3* conditional mice were F: 5’-GAGATGCTGATTTCACTGTCTAGC-3’ and R: 5’-CTCTTATCACTGGCTGAAACTACA-3’. PCR primers for the *Cre recombinase* transgene used were Forward: 5’-GAACGCACTGATTTCGACCA-3’ Reverse: 5’-GCTAACCAGCGTTTTCGTTC-3. Skeletal myofiber tamoxifen-inducible mice were purchased from Jackson Labs (Bar Harbor, ME) Human Skeletal-Actin-MerCreMer (*HSA-MCM*) mice (Jackson Labs; Bar Harbor, ME; stock# 025750) and WT (*C57BL/6J*; stock# 000664) were maintained in our animal colony under pathogen-free standard housing conditions. *Dock3* ubiquitous KO mice (Jackson Labs; stock# 033736) were originally obtained from the laboratory of Dr. David Shubert (Salk Institute) and have been previously described (10). The *mdx*^*5cv*^ (Jackson Labs; stock# 002379) mice were originally obtained from Jackson Labs. All mice were maintained on the *C57BL/6J* strain background. All mouse strains were maintained under standard housing and feeding conditions with the University of Alabama at Birmingham Animal Resources Facility under pathogen-free, sterile conditions under animal protocol number 21393. Mice were all fed a diet consisting of the Teklad Global Rodent Diets (Envigo; Indianapolis, IN) with *ad libitum* access to food and water.

### GLUT4-transfection

WT and *Dock3* KO primary mouse muscle cells were harvested from the ubiquitous *Dock3* KO mice as previously described(Chen, Peto et al. 2009). Primary mouse myoblasts were grown in Skeletal Muscle Cell Growth Medium (Promocell Cat# C-23060; Heidelberg, Germany) with 20% FBS (ThermoFisher Scientific; Waltham, MA; Cat# 16140071), and incubated at 37°C using a standard primary muscle cell isolation protocol(Gharaibeh, Lu et al. 2008). Muscle cells were plated in a 6-well gelatin coated plate at 50,000 cells/well. Muscle cells were then transfected with a pLenti-myc-GLUT4-mCherry (Addgene; Watertown, MA; stock # 64049) for 48 hours and the GLUT4 localization assay was performed as previously described(Lim, Bi et al. 2015).

### Mouse activity tracking

Mouse activity locomotor measurements were performed as previously described(Reid, Wang et al. 2020). Twenty-four hours prior to experiment termination and tissue harvest, mice were analyzed for locomotive activity using the Ethovision XT software platform (Noldus; Leesburg, VA) with isolated individual chambers that recorded motion from mouse head to tail. Mice were acclimated to the room and open-field chambers one day prior to activity and were given a five minute additional adaptation period prior to activity recording. Mouse activity was recorded for six minutes with no external stimulation.

### Myofiber diameter calculations

The cross-sectional area (CSA) of the myofibers within the skeletal muscle sections was calculated by quantifying the myofiber areas using a previously described protocol(Mula, Lee et al. 2013). Approximately 600 TA myofibers were counted and CSA (µm^2^) was measured via several overlapping H&E microscopy images of each section and quantified using Fiji software(Schindelin, Arganda-Carreras et al. 2012).

### DEXA Quantitative Magnetic Resonance (QMR) imaging

Evaluation of body composition comprising of both fat and lean tissue mass *in vivo* was performed on 4-month-old male *Dock3*^*fl/fl*^ and *Dock3* mKO mice (10 mice/genotype) using the EchoMRI™ 3-in-1 composition analyzer (software version 2016, Echo Medical; Houston, TX). Individual fat and lean mass measurements were recorded in grams (g) and were analyzed using student’s t-test, two-tailed between *Dock3*^*fl/fl*^ and *Dock3* mKO mice.

### Cardiotoxin-induced skeletal muscle injury

*Dock3* mKO and *Dock3*^*fl/fl*^ mice were injected in their TA skeletal muscles with 40 µl of cardiotoxin (MilliporeSigma; Cat# 217503) at a 10 µM concentration. The contralateral TA muscle was used as a sham control injection with 1x phosphate-buffered saline (ThermoFisher Scientific; Cat# 10010049). Seven days following injections, mice were euthanized, and their TA skeletal muscles were slow-frozen in Scigen TissuePlus O.C.T. Compound (Fisher Scientific; Hampton, NH Cat# 23-730-571) for histological analysis and snap frozen in liquid nitrogen for molecular analysis.

### Immunofluorescence and immunohistochemistry

Mouse skeletal muscles were cryo-frozen in Scigen TissuePlus O.C.T. Compound using an isopentane (FisherScientific; Cat# AC397221000) and liquid nitrogen bath as unfixed tissues. Blocks were later cut on a cryostat into 7-10 µm sections and placed on Fisherbrand Tissue Path Superfrost Plus Gold slides (Fisher Scientific; Cat# FT4981gplus). H&E staining was performed as previously described(Beedle 2016). For immunofluorescent staining, slides were blocked for one hour in 10% goat serum and incubated for one hour at room temperature using a M.O.M kit (Vector Labs Cat# BMK-2202; Newark, CA).

### Western blotting

Protein lysates were obtained by homogenizing tissues in M-PER lysis buffer (ThermoFisher; Cat# 78501) with 1x Complete Mini EDTA-free protease inhibitor cocktail tablets (Roche Applied Sciences; Cat# 04693159001; Penzburg Germany). Protein lysates were quantified using a Pierce BCA Protein Assay Kit (ThermoFisher Cat# 23225). Unless stated otherwise, 50 µg of whole protein lysate was used for all immunoblots and resolved on 4-20% Mini-PROTEAN TGX Precast Protein gels (BioRad; Cat# 4561094). Protein samples were transferred to 0.2 µm PDVF membranes (ThermoFisher; Cat# LC2002), blocked in 0.1x TBS-Tween in 5% BSA for one hour, and then gently incubated overnight with primary antibody on a rocker at 4°C. Membranes were washed in 0.1% TBS-tween four times at 10-minute intervals before being incubated with secondary antibodies (either mouse or rabbit IgG) conjugated to HRP for one hour at room temperature with gentle agitation. Following another three washes for 15 minute intervals at room temperature, membranes were then treated with RapidStep ECL Reagent (MilliporeSigma; Cat# 345818-100 ml).

### Real-time quantitative PCR

Total RNA was extracted using a miRVana (ThermoFisher, Cat# AM1560) kit while following the manufacturer’s protocol. One microgram of total RNA was reverse transcribed using the Taqman Reverse Transcription kit (Applied Biosystems; Cat# N8080234; Waltham, MA) following the manufacturer’s protocol. TaqMan assay probes were all purchased from ThermoFisher corresponding to each individual transcript. Quantitative PCR (qPCR) TaqMan reactions were performed using TaqMan Universal PCR Master Mix (Applied Biosystems; Cat# 4304437). Relative expression values were calculated using the manufacturer’s software and further confirmed using the 2^-ΔΔ^Ct method.

### Glucose and insulin tolerance tests

Mice were fasted for eight hours prior to afternoon administration of a bolus of D-glucose (MilliporeSigma; Cat# G8270). Mice were given an intraperitoneal injection at a concentration of 3 mg/gram of mouse bodyweight. Blood glucose was measured on a commercially obtained glucometer (Nipro Diagnostics Inc.; Southampton, UK) using 10 µl of whole serum from tail bleeds. For the insulin tolerance tests, the mice were fasted for five hours prior to afternoon administration of a bolus of human insulin (MilliporeSigma; Cat# 1342106). Mice were given an intraperitoneal injection at a concentration of 3 mg/gram of mouse bodyweight. Blood glucose was measured on a commercially obtained glucometer using 10 µl of whole serum from tail bleeds.

### Yeast-2-Hybrid

The GAL4-based yeast two hybrid system was used to detect the interaction between recombinant DOCK3 and SORBS1 domains. The bait and prey are expressed as fusion domain constructs to the GAL4 DNA binding domain and GAL4 activation plasmids. Inoculations were then transferred to a 500 mL flask containing 300 mL yeast peptone dextrose (YPD) broth (ThermoFisher Scientific; Cat# A1374501) and incubated at 30°C for 16-18 hours with shaking at 230 rpm. Cultures were incubated at 30°C for 16-18 hours with shaking at 230 rpm in an overnight culture flask containing 300 ml of YPD. Cultures were harvested in 50 ml tubes and centrifuged at 1000 x g for five minutes at room temperature. Cell pellets were resuspended in distilled water and again centrifuged at 1000 x g for five minutes. Pellets were then re-suspended in 1.5 ml freshly prepared, sterile 1X TE/1X LiAc solution. Approximately 0.1 µg of plasmid DNA and 0.1 mg of carrier DNA was added to a 1.5 mL tube and mixed. Approximately 0.1 ml of yeast competent cells were then added to each tube and vortexed until well mixed, heat shocked for five minutes in a 42°C water bath, and chilled on ice for 2 minutes. Yeast cultures were then centrifuged for five seconds at 12,000 x g and resuspended in 0.5 µL sterile TE buffer. The cells were plated at 100 µL each on SD/-LEU/-Trp selective transformant agar plates and incubated at 30°C until colonies appeared the next morning.

### Co-immunoprecipitation (co-IP)

Protein constructs were expressed in HEK293T cells using Lipofectamine 2000-mediated (Invitrogen, Catalog #11668030; Waltham, MA) plasmid transfection. Expression constructs were subcloned into Vitality hrGFP mammalian expression vectors (Agilent Technologies; Santa Clara, CA; Cat# 240031 and #240032) using standard PCR cloning techniques. HEK293T cells were collected two days post-transfection and lysed in lysis buffer that contained 50 mM Tris-HCl (pH 7.4), 150 mM NaCl, 1 mM EDTA, 1% Triton X-100, and 1:100 Protease/Phosphatase Inhibitor Cocktail (Cell Signaling Technology; Danvers, MA). Cells were then homogenized using an Omni Bead Rupter 12 (Perkin Elmer; Kennesaw, GA). Protein lysates were then incubated on ice for thirty minutes. Lysates were spun down at 10000 x g for ten minutes, and the supernatant was collected for co-IP. Protein levels were quantified using the BCA Kit and normalized (Pierce Protein Biology, Rockford, IL, USA). Approximately 5% of total protein lysate was set aside as the input fraction. Laemmli Buffer plus β-mercaptoethanol was then added to these samples and one mg of total protein lysate was used per co-IP reaction. Approximately 0.5 mg of mouse IgG control (ThermoFisher, Catalog # MA1-213) was used for the control reaction. Co-IP reactions were rotated overnight at 4°C with 100 µl of SureBeads Protein G Magnetic Beads (BioRad; Catalog# 1614013; Hercules, CA). The bead lysates were washed five times in the co-IP buffer using a DynaMag Magnet (ThermoFisher) to pull down the complexes. After this, Laemmli Buffer plus β-mercaptoethanol was added to the beads, which were boiled for five minutes at 100°C. All co-IP reactions were probed using standard western immunoblotting techniques described above. The rabbit DOCK3 (ThermoFisher; Cat# PIPA5100485) and mouse SORBS1 (Sigma-Aldrich; Catalog #SAB4200599; St. Louis, MO) antibodies were used for verifying immunoprecipitation reactions via western immunoblotting. Anti-FLAG M2 magnetic beads (MilliporeSigma; Catalog #M8823) and anti-FLAG M2 monoclonal antibody (MilliporeSigma; Catalog #F1804) were used for co-IP and western immunoblotting reactions. A µMACS HA magnetic bead isolation kit (Miltenyi Biotec; Catalog# 130-091-122) and anti-HA rabbit monoclonal (GenScript; Catalog #A01963) were also used for co-IP and western immunoblotting reactions.

### GST pulldown assay

Recombinant SORBS1 protein (Abcam; Cambridge, UK) was incubated with recombinant GST-DOCK3-PXXP or GST alone plasmids (constructs cloned into pGEX-6P-1 plasmid; GE Healthcare; Chicago, IL) in GST reaction buffer (250 mM Tris-HCl at pH 7.4, 500 mM NaCl, 25 mM MgCl_2_, 5 mM dithiothreitol, 0.5 mM EGTA and 20 mM freshly prepared ATP) for one hour at 4°C on a rotator. Pierce Glutathione Magnetic Agarose Beads (ThermoFisher; Cat# 78602) were then suspended in the GST reaction buffer and added to the reaction mixture for one hour at 4°C with gentle rotation. The beads were then washed four times in reaction buffer using a DynaMag magnet. Laemmli Buffer plus β-mercaptoethanol was added to these samples, which were then boiled for five minutes at 100°C. GST pulldown was verified via immunoblot against the GST epitope (anti-GST; rabbit polyclonal; Abcam; Cat# ab9085).

### Muscle physiological function assays

EDL muscles were dissected from anesthetized mice and studied in a phosphate buffer equilibrated with 95% O_2_, 5% CO_2_ (35 °C). Contractions were produced using a 150 ms, supramaximal stimulus train (200 µs pulses) with the muscle held at its optimal length (L_o_) for tetanic tension. Force was normalized to physiological cross-sectional area as previously described (Huntoon et al. 2018). Each muscle was studied at stimulation frequencies ranging from 30 to 300 Hz (peak force). Fixed-end force values were expressed relative to peak force and fit by a sigmoid curve as previously described(Huntoon, Widrick et al. 2018). Changes in the relationships were evaluated by differences in the inflection point (K, measured in Hz) and slope (H, unitless). Muscles were then subjected to a high active strain protocol consisting of the following sequence: one fix-end trial, 5 lengthening (eccentric) trials, and two fixed-end trials. The fixed-end trials were as described above. The lengthening trials consisted of an initial fixed-end contraction that allowed the muscle to rise to peak force (100 ms duration), followed by a constant velocity stretch at 4 fiber lengths/s (50 mms duration) to a final length of 120% L_o_. For the high-strain protocol, force was evaluated at 95 ms of stimulation for both fixed-end and lengthening trials.

### Statistical analyses

Unless otherwise described, a two-tailed student’s t-test was performed for all single comparisons and either a one-way or two-way analysis of variance (ANOVA) with Tukey’s post-hoc honest significant difference (HSD) was performed for all multiple comparisons. GraphPad Prism version 9 software (Graphpad Software; San Diego, CA) was used for all statistical analyses. An *a priori* hypothesis of *p< 0.05, **p<0.01, ***p<0.001, and ****p< 0.0001 was used for all reported data analyses. All graphs were represented as mean +/-SEM.

## Results

### Generation of a muscle-specific Dock3 knockout mouse

As DOCK3 protein is expressed both within the motor neuron and in the skeletal muscle, we generated a conditional mouse model to differentiate the role of the *Dock3* gene exclusively in skeletal muscle. We investigated the specific function of *Dock3* in the myofiber by evaluating *Dock3*-deficient mice in which exons 8 and 9 of the *Dock3* gene locus is flanked with loxP sites in the intronic regions (**Figure 1A**). Upon mating with the mouse model expressing tamoxifen-inducible *Cre recombinase* driven by the human-skeletal actin promoter (*HSA-MerCre-Mer*; *HSA-MCM*), the mice will conditionally ablate *Dock3* expression upon tamoxifen administration in the skeletal myofibers **(Figure 1A)**. Genotyping and western blot analyses of *Dock3* expression in the tissue extracts of brain and the tibialis anterior (TA) from control and *Dock3* ubiquitous KO mice confirmed the ablation of *Dock3* from the myofiber. The *Dock3*^*fl/fl*^*:HSA-MerCreMer* (henceforth referred to as *Dock3* mKO) mice showed the deletion of *Dock3* expression in tissue extracts from the TA lysates, but not the brain, confirming the tissue-specific deletion of *Dock3* from the myofiber (**Figures 1B, 1D, and 1E**).

**Figure 1.**
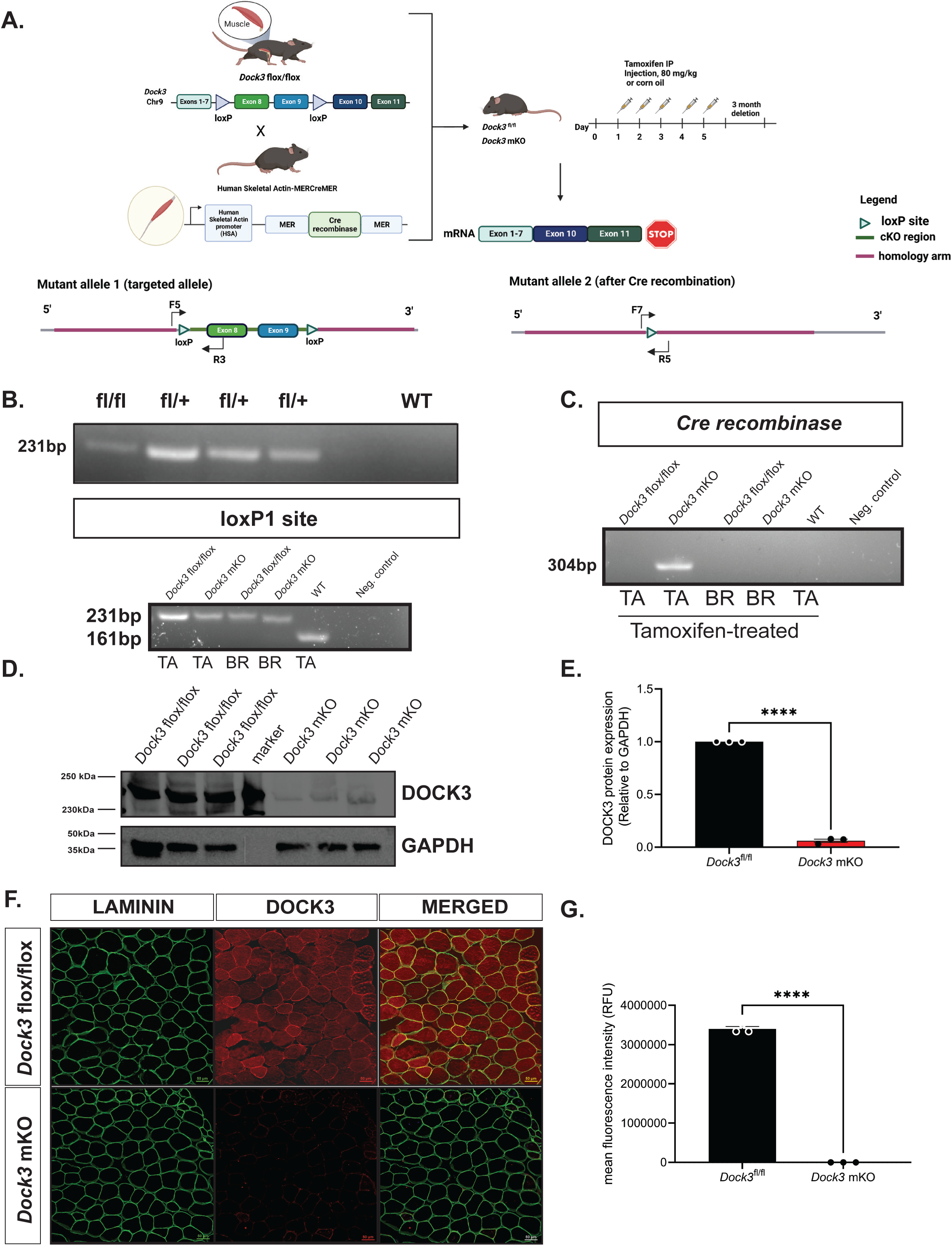
Generation and validation of muscle-specific *Dock3* conditional knockout mice. **A**. Generation of *Dock3* mKO schematic. *Dock3*^*fl/fl*^ mice containing two loxP sites flanking *Dock3* exon 8 and 9 were mated with human-skeletal-actin (*HSA*)-*MerCreMer* mouse line to generate the *Dock3* mKO mice. When administered tamoxifen (80 mg/kg) over five consecutive days this induces a frameshift mutation resulting in a premature stop codon. **B**. PCR genotyping agarose gel of *Dock3* heterozygous fl/+ alleles to produce homozygous flox/flox *Dock3* in skeletal muscle. **C**. PCR genotyping agarose gel identifying loxP1 site (231 bp) and *Cre* recombinase (304 bp) in *Dock3* flox/flox and WT mice (161 bp) mice in both tibialis anterior (TA) and whole brain lysates (BR). **D**. Western blot of *Dock3* mKO mice indicating a reduction of protein as a result of ablation of *Dock3*. **E**. Quantification of DOCK3 protein normalized to GAPDH loading control. **F**. Immunofluorescent staining of adult TA muscles from the *Dock3*^*fl/fl*^ and *Dock3* mKO mice for LAMININ, DOCK3, and the merged image. Scale bar = 50 µm. **G**. Quantification of mean fluorescence intensity (RFUs) in the *Dock3*^*fl/fl*^ and *Dock3* mKO mice. Significance shown as ****p< 0.0001.

### Dock3 mKO mice have disrupted skeletal muscle histology and locomotor activity

To evaluate the consequences of *Dock3* skeletal muscle ablation, we first analyzed the muscle architecture and morphology of isolated TA muscle fibers of 4-month-old *Dock3* mKO mice compared to *Dock3*^*fl/fl*^ controls **(Figure 2A)**. We observed a decrease in myofiber cross-sectional area (CSA) and noted smaller myofibers grouped together throughout the *Dock3* mKO muscles (**Figure 2B**). We did not observe a change in centralized myonuclei. However, we did observe muscle fiber atrophy reflected by increased frequency of smaller myofibers in *Dock3* mKO mice compared to *Dock3*^*fl/fl*^ controls. We sought to characterize how the disruption of *Dock3* in the skeletal muscle would impact overall locomotive function by using open field activity tracking to record activity levels in adult mice (**Figure 3A**). *Dock3* mKO mice demonstrated significantly decreased distance traveled and average velocity compared with controls, indicating a reduction in basal locomotor function (**Figures 3B-D**). These findings are consistent with a decrease in locomotor function previously observed in adult ubiquitous *Dock3* KO mice. Interestingly, when we conducted several functional assays on extensor digitorium longus (EDL) muscles isolated from *Dock3* mKO and *Dock3*^*fl/fl*^ mice we found no significant changes in the muscle’s contractile properties (**Figures 3E-3J**). This included the relationship between stimulus frequency and force (**Figures 3E-G**), absolute peak force (**Figure 3G**), force normalized to the muscle’s physiological cross-sectional area (**Figure 3I**), and in the muscles resistance to eccentric contractions (**Figures 3J and 3K**). Therefore, we concluded that loss of *Dock3* in the myofiber reduces basal activity independent of undermining the overall contractility and structural integrity of the skeletal muscle.

**Figure 2:**
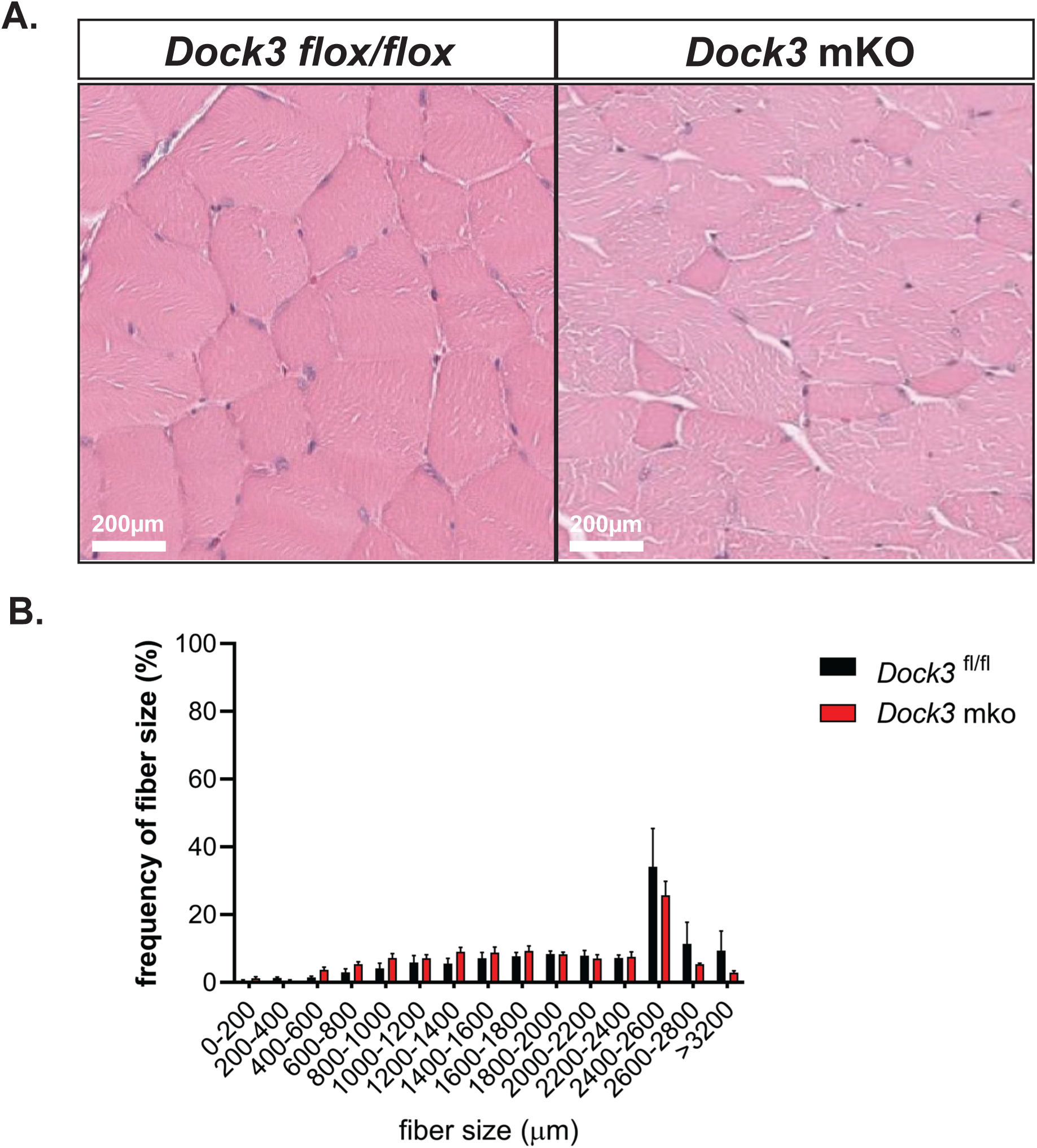
Muscle-specific loss of *Dock3* results in a smaller myofiber sizes. **A**. Hematoxylin and eosin (H&E) stainings of TA muscles from *Dock3*^*fl/fl*^ vs. *Dock3* mKO. Scale bar = 200 µm. **B**. Quantification of myofiber diameters in *Dock3*^fl/fl^ vs. *Dock3* mKO. Cross-sectional area shown as frequency of fiber sizes over fiber size (µm^2^).

**Figure 3.**
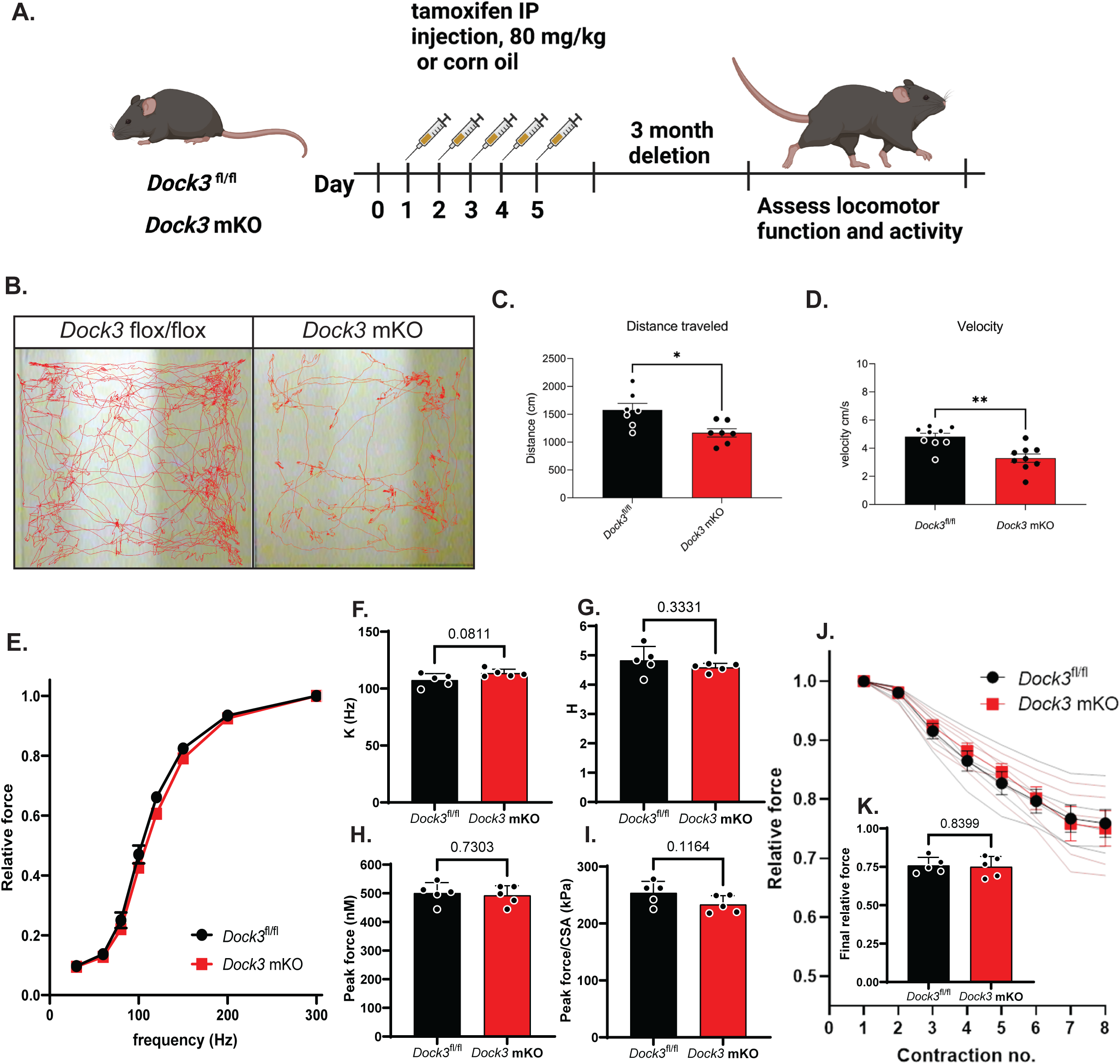
Loss of muscle *Dock3* reduces locomotor activity independent of physiological force. **A**. Schematic showing tamoxifen regimen for *Dock3*^*fl/fl*^ and *Dock3* mKO. Mice were administered an intraperitoneal injection of tamoxifen (80 mg/kg) for five consecutive days followed by a three month washout period before assessing locomotor function. **B**. Activity tracking traces in *Dock3*^*fl/fl*^ vs. *Dock3* mKO mice. **C**. Quantification of total distance traveled (cm) n = 9 mice/cohort, **D**. Quantification of mouse velocity, n = 9 mice/cohort, **E**. Force-frequency relationship of EDL muscles from *Dock3* mKO vs. *Dock3*^*fl/fl*^ EDL mice, n = 5 mice/cohort. **F**. Inflection point of the force-frequency relationship (K) of EDL muscles from *Dock3* mKO vs. *Dock3*^*fl/fl*^ mice n = 5 mice/cohort. **G**. Slope of the force-frequency relationship (H) of EDL muscles from *Dock3* mKO vs. *Dock3*^*fl/fl*^ mice n = 5 mice/cohort. **H**. Peak force of EDL muscles from *Dock3* mKO vs. *Dock3*^*fl/fl*^ mice n = 5 mice/cohort. **I**. Peak force per physiological cross-sectional area (CSA) of EDL muscles from *Dock3* mKO vs. *Dock3*^*fl/fl*^ mice, n = 5 mice/cohort. **J**. Relative isometric force measured during the eccentric contraction protocol for EDL muscles of *Dock3* mKO vs. *Dock3*^*fl/fl*^ mice, n = 5 mice/cohort. **K**. Relative force at the conclusion of the eccentric contraction protocol for EDL muscles of *Dock3* mKO vs. *Dock3*^*fl/fl*^ mice, n = 5 mice/cohort. The following p-values of significance were stated: *p < 0.001, **p < 0.01, and ns = not significant.

### Loss of Dock3 at the myofiber inhibits myogenic regeneration after cardiotoxin injury

Previously, we isolated primary myoblasts isolated from *Dock3* KO muscle which exhibited impaired regeneration and fusion. We sought to determine if this phenomenon was recapitulated in our *Dock3* mKO mice and the degree to which muscle regeneration is impacted by the loss of muscle DOCK3 expression **(Figure 4A)**. We performed an intramuscular injection of cardiotoxin in our *Dock3* mKO mice to induce a skeletal muscle injury into the right TA muscle while using the left as a contralateral control receiving a sham injection of phosphate buffered saline (PBS) to evaluate the role of DOCK3 in muscle regeneration. Mice were sacrificed at 7 days post-injury and evaluated via histological analysis with hematoxylin and eosin (H&E) and Masson’s trichrome to assess myofiber cross-sectional area, myonuclei position, and fibrosis within the muscle. We observed that the *Dock3* mKO mice had increased fibrosis when compared to the control *Dock3*^*fl/fl*^ **(Figures 4B and 4C)**. We also quantified increased levels of centralized myonuclei in the *Dock3* mKO mice (**Figure 4C**). We repeated the study for 14 days post cardiotoxin TA muscle injury and observed similarly impaired muscle regeneration in the *Dock3* mKO mice as seen in the *Dock3* global KO mice (**Figure 4D**). These findings were consistent with the high levels of centralized myonuclei and fibrotic areas observed, indicating a delay in regeneration in the skeletal muscle of *Dock3* mKO mice and emphasizing the importance of DOCK3 in skeletal muscle.

**Figure 4.**
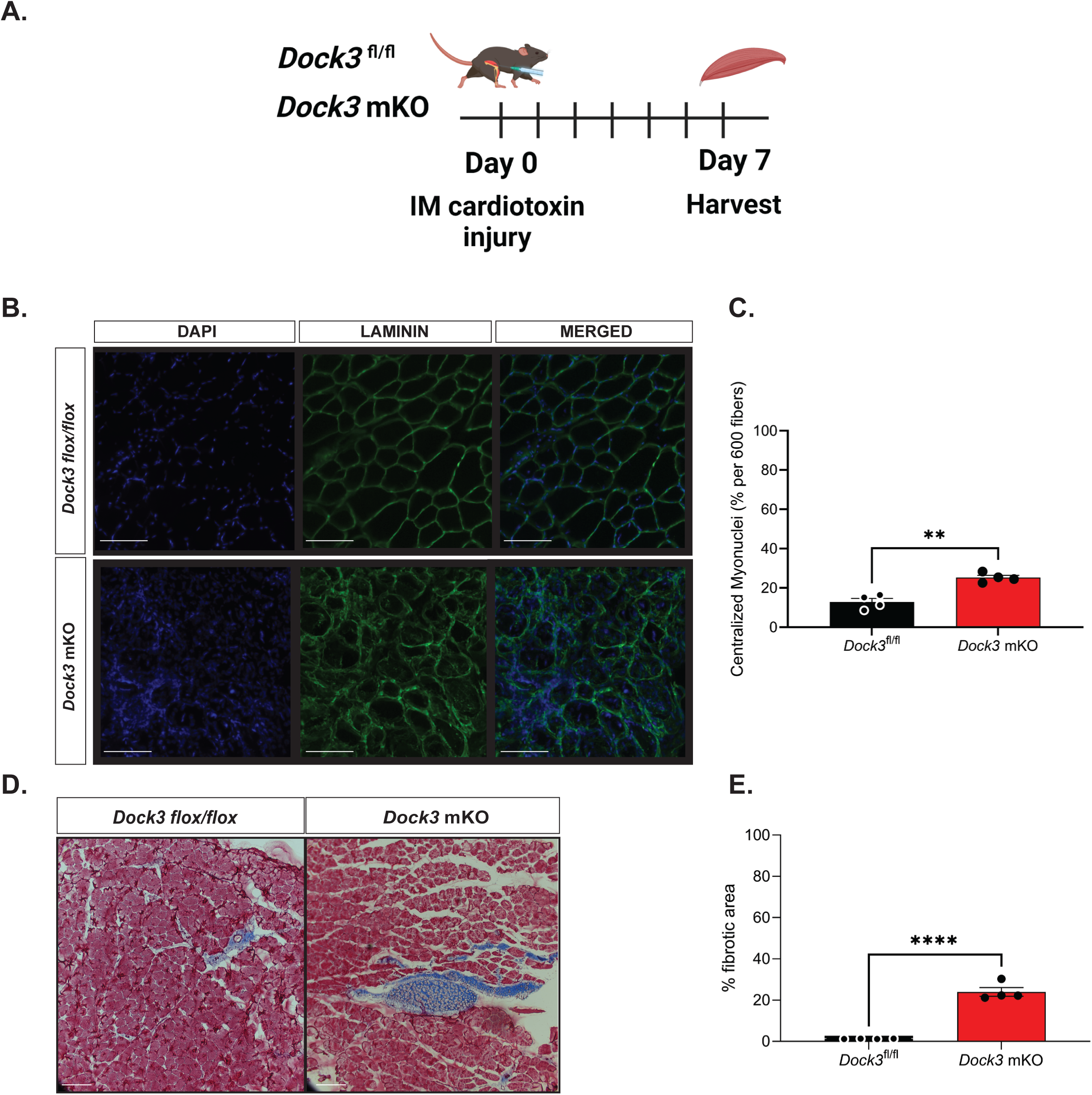
*Dock3* mKO mice show impaired skeletal muscle regeneration following injury. **A**Schematic of cardiotoxin induced skeletal muscle TA injury. *Dock3* mKO mice and *Dock3*^*fl/fl*^ mice were administered with an intramuscular injection of 10 µM of cardiotoxin at Day 0 and sacrificed and harvested day 7 post-injury. **B**. Cross-section of injured tibialis anterior stained with immunofluorescent antibody against LAMININ (green), DAPI (blue), and the merged image. Scale bar = 200 µm. **C**. Quantification of immunofluorescent images from (**B**) analyzing % centralized myonuclei per 600 fibers. **p < 0 .001, n = 4 mice/cohort. **D**. Masson’s trichrome histochemical analysis of injured TA in *Dock3*^*fl/fl*^ vs. *Dock3* mKO 7 days post-injury. **E**. Quantification of histochemical images from (**D**) with analysis of percent (%) fibrotic area in injured TA of *Dock3*^*fl/fl*^ vs. *Dock3* mKO 7 days post-injury, n = 4 mice/cohort, p < 0.0001. Scale bar = 200 µm.

### Adult Dock3 mKO mice have abnormal skeletal muscle mass and metabolism

We previously demonstrated that *Dock3* ubiquitous KO mice were glucose intolerant and had decreased weights due to decreased muscle mass. Thus, we sought to understand if the loss of *Dock3* in the skeletal muscle would impact whole-body metabolism. Quantitative magnetic resonance (QMR) imagining of adult *Dock3* mKO mice revealed increased body weight compared to *Dock3*^*fl/fl*^ aged-matched controls **(Figure 5A)**. Conversely, *Dock3* mKO mice had significantly increased fat mass compared to *Dock3*^*fl/fl*^ aged-matched controls (**Figure 5B)**. No detectable changes in skeletal muscle lean mass were observed in the *Dock3* mKO mice **(Figure 5C)**. Being that DOCK3 is known to activate Rho GTPases such as RAC1, a critical regulator of insulin and glucose signaling pathways in skeletal muscle. We measured the ability of the *Dock3* mKO mice to respond to a glucose challenge via a glucose tolerance test (GTT). GTT tests revealed no significant changes in glucose processing in the muscle **(Figure 5D)**. However, insulin tolerance tests (ITT) conducted on *Dock3* mKO mice revealed whole body hyperglycemia and insulin resistance **(Figure 5E)**. We analyzed the role of DOCK3 in glucose processing within the muscle by isolating primary *Dock3* KO myoblasts and infecting with lentiviral GLUT4-RFP(Wang, Khayat et al. 1998). Upon insulin stimulation, we observed reduced GLUT4 translocation in the *Dock3* KO myoblasts, which supports a defect in glucose uptake and/or processing within the skeletal muscle (**Figure 5F**). These findings reveal DOCK3 to be a critical regulator of metabolism in the skeletal muscle and that loss of DOCK3 expression in the myofiber undermines important metabolic functioning and insulin processing in the skeletal muscle.

**Figure 5.**
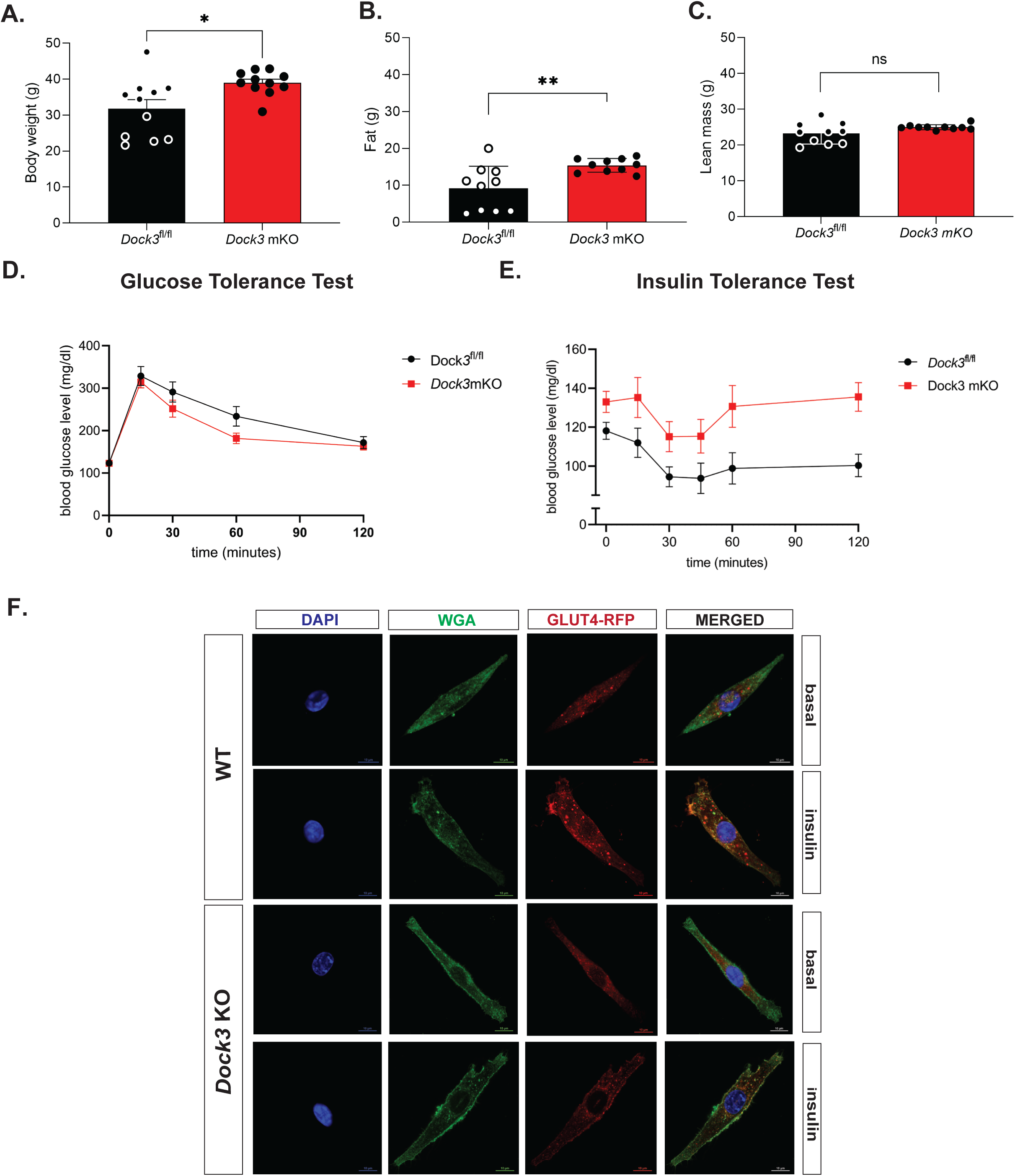
*Dock3* mKO mice show increased body mass and whole body hyperglycemia. **A**. Quantitative magnetic resonance imaging indicated body weight differences between *Dock3*^*fl/fl*^ and *Dock3* mKO mice, n = 10 mice/cohort, *p < 0.01. **B**. Quantitative magnetic resonance imaging indicated differences in fat mass between *Dock3*^*fl/fl*^ and *Dock3* mKO mice, n = 10 mice/cohort, *p < 0.001. **C**. Quantitative magnetic resonance imaging indicating differences in lean mass between *Dock3*^*fl/fl*^ and *Dock3* mKO mice, n = 10 mice/cohort, ns = not significant. **D**. Glucose Tolerance Test in *Dock3*^*fl/fl*^ and *Dock3* mKO mice shown. Serum blood glucose level (mg/dl) measured over time (minutes). **E**. Insulin Tolerance Test in *Dock3*^*fl/fl*^ vs. *Dock3* mKO mice. n = 8 mice/cohort. Serum blood glucose level (mg/dl) measured over time (minutes). **F**. WT and *Dock3* KO myoblasts transfected with HA-GLUT-RFP. Wheat germ agglutinin stained membranes (green), GLUT4 (RFP), and nuclei are stained with DAPI. Scale bar = 10 µm.

### DOCK3 interacts with insulin signaling protein, Sorbin and SH3 domain-containing 1 (SORBS1)

Due to the increased fat mass, body weight, and hyperglycemia observed in the *Dock3* mKO mice, we explored what potential protein-protein interactions DOCK3 may be involved with regarding glucose uptake. We conducted a yeast two-hybrid neuromuscular cDNA library screen using the C-terminal domain of human DOCK3 protein to identify novel DOCK3 protein interactions (**Figure 6A**). We identified the insulin adaptor protein, SORBS1 as directly interacting with the C-terminal domain of DOCK3 and confirmed the interaction via secondary yeast amino acid growth selection confirmation (**Figure 6B and Figure 6C**). SORBS1, also called Cbl-Associated Protein (CAP), is a known insulin adaptor protein whose subcellular localization is essential to downstream insulin signaling events and has been implicated as a secondary signaling pathway critical to insulin-mediated glucose uptake(Baumann, Ribon et al. 2000). To determine which domains of each protein were critical to their interaction, we conducted a GST-pulldown assay in HEK293T cells overexpressing DOCK3 and SORBS1. The proline rich motif (PXXP) of DOCK3 and the SH3 domains of SORBS1 were identified as the main sites of the DOCK3-SORBS1 protein interaction (**Figures 6D-6F**). Following these results, we sought to map out which functional domains were critical to the DOCK3-SORBS1 interaction. Overexpression constructs containing full length and deletion of key conserved protein functional domains of DOCK3 and constructs deleting each of the SH3 domains of SORBS1 were generated **(Figure 7A)**. Co-immunoprecipitation confirmed that all three SH3 domains on SORBS1 were essential for the DOCK3-SORBS1 interaction **(Figure 7B)**. This protein-protein interaction between DOCK3 and SORBS1 was further validated in human primary myotubes **(Figures 7C-7D)**. These results identified the DOCK3-SORBS1 interaction as a potential novel source of metabolic regulation that may modulate glucose and insulin signaling in skeletal muscle.

**Figure 6.**
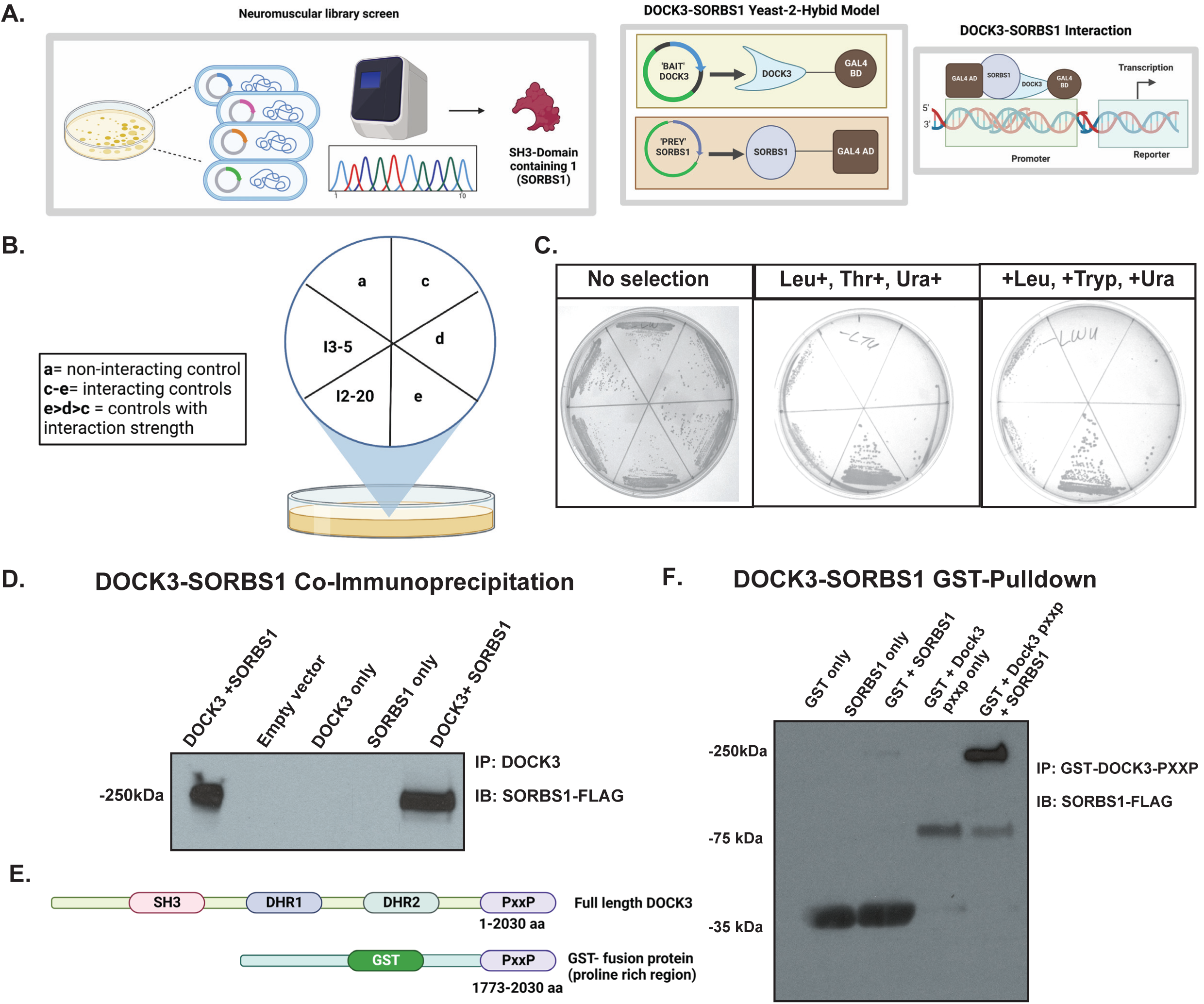
DOCK3 interacts with insulin signaling protein, SH3 domain-containing 1 (SORBS1). **A**. DOCK3-SORBS1 yeast-2-hybrid neuromuscular cDNA screening library strategy. Selection guide of DOCK3-SORBS1 yeast-2-hybrid amino acid selection. **C**. Positive interaction of DOCK3-C-terminus and SORBS1 cDNA shown with yeast selective growth. **D**. DOCK3-SORBS1 co-Immunoprecipitation in HEK293T cells. Immunoprecipitation (IP) performed with DOCK3-GST and immunoblotting (IB) against SORBS-FLAG. **E**. GST-Pulldown domain constructs shown indicating domains of full-length Dock3 and GST tagged PXXP motif. **F**. DOCK3-SORBS1 GST-Pulldown immunoblots showing the interaction between recombinant DOCK3 and SORBS1 directly interacting.

**Figure 7.**
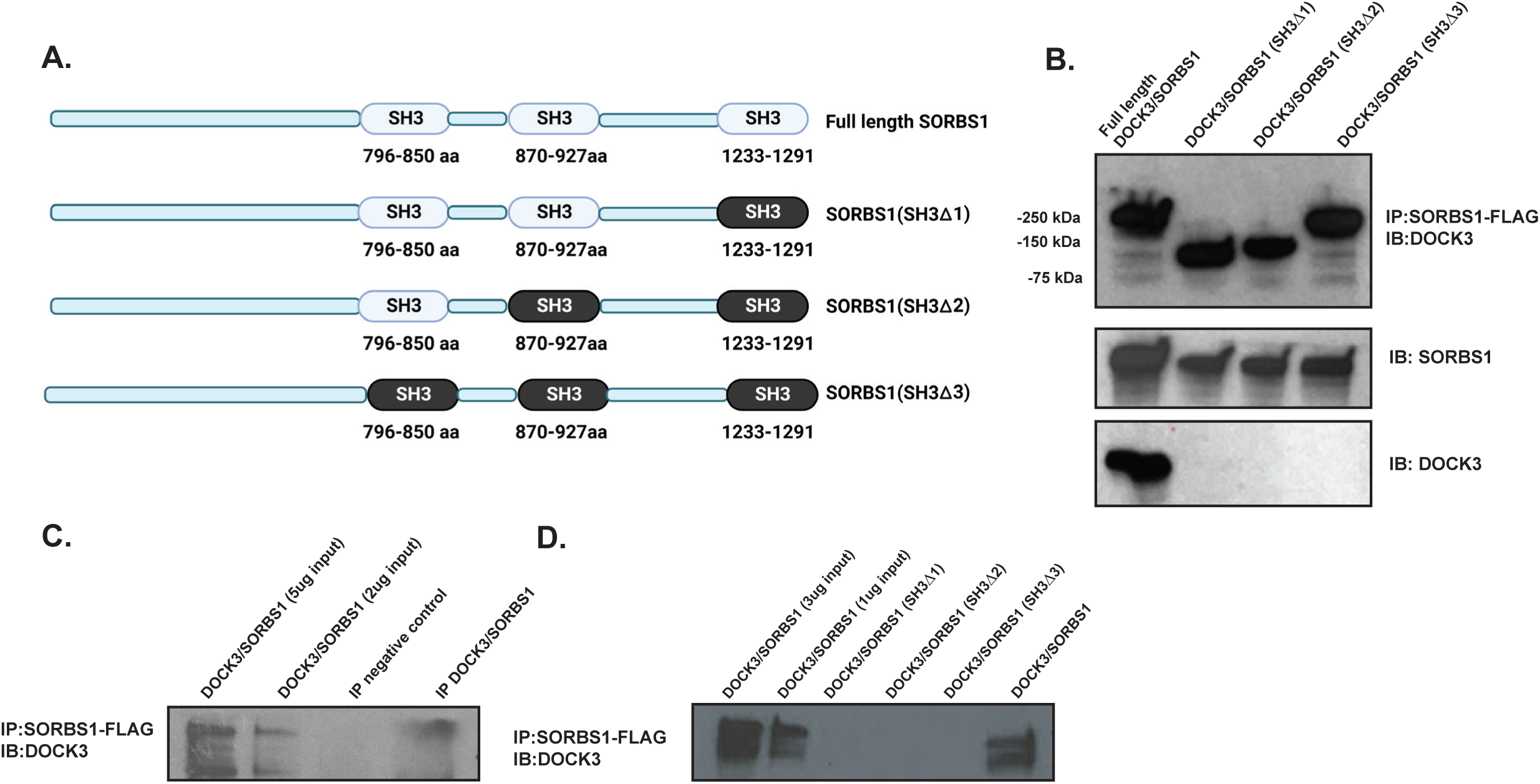
DOCK3 interacts with SORBS1 via binding to the SORBS1 SH3 domains. **A**. Schematic showing the SORBS1 deletion constructs indicating full length SORBS1, and deletion domains across each SH3 domain. **B**. The co-IP indicating the interaction and expression of each deletion construct in HEK293T cells. **C**. The co-IP of DOCK3-SORBS1 in human primary myoblasts. Immunoblot (DOCK3 rabbit polyclonal antibody) with the co-immunoprecipitation (SORBS1-FLAG; FLAG mouse monoclonal antibody) **D**. The co-IP of SORBS1 deletion constructs containing deletions of each of the SH3 domains in SORBS1 showing the requirements for each in binding to DOCK3.

## Discussion

DOCK3 is a guanine-nucleotide exchange factor whose downstream activation of Rho GTPases impacts a number of pathways that influence cell migration, insulin signaling, and pathways regulating muscle mass(Bryan, Li et al. 2005, Chiu, Jensen et al. 2011). Our previous work identified DOCK3 as an important biomarker and dosage-sensitive regulator of Duchenne muscular dystrophy(Reid, Wang et al. 2020). Here, we identified the muscle-specific role of DOCK3 in the myofiber, apart from its role in the motor neuron, and how its expression is expression is critical to normal muscle function, regeneration and glucose processing within the muscle. Furthermore, using a yeast-two-hybrid screen we identified a novel protein-protein interaction with SORBS1, a key glucose and insulin signaling factor that may yield clues into DOCK3’s regulation of skeletal muscle metabolic function via glucose and insulin signaling pathways.

Additional questions remain as DOCK3 has been shown to interact with RAC1, another key regulator of glucose processing in skeletal muscle which may also explain our observed phenotypes in the *Dock3* mKO mice(Sylow, Jensen et al. 2013, Li, Mi et al. 2016, Raun, Ali et al. 2018). Our observation of impaired skeletal muscle regeneration following cardiotoxin injury in the *Dock3* mKO mice suggests that DOCK3 may play roles in muscle regeneration even though it is not expressed in the muscle satellite or stem cells. DOCK3 has been shown to play key roles in cell migration, actin polymerization, and regulates key signaling pathways such as WAVE which may explain the observed impaired regeneration in the *Dock3* mKO mice(Namekata, Harada et al. 2010). The observation of *Dock3* mKO mice having whole-body hyperglycemia and insulin resistance could be a result of the disruption of the DOCK3-SORBS1 interaction. SORBS1 is part of a small family of adaptor proteins that is known to regulate numerous cellular processes including cell adhesion, cytoskeletal formation, and is required for insulin-stimulated glucose transport(Mandai, Nakanishi et al. 1999). A number of studies suggest that genetic variations in SORBS1 could be associated with human disorders such as obesity, diabetes, and insulin resistance(Baumann, Ribon et al. 2000, Lesniewski, Hosch et al. 2007, Chang, Wang et al. 2018). *Dock3* mKO mice showed significantly increased fat mass and body mass, without any impact on lean mass. The novel interaction between DOCK3 and SORBS1 implies that DOCK3 plays a significant metabolic role in the muscle, specifically involving regulation of insulin-mediated glucose uptake. Further studies are warranted to dissect DOCK3’s additional roles in other lineages using a conditional approach.

## Author Contributions

A. Samani, M. Karuppasamy, K. English, C. Siler, Y. Wang, and J. Widrick all performed experiments related to the project and analyzed the data. A.Samani, M. Karuppasamy, K. English, and M. Alexander all analyzed the data and wrote, edited, and revised the manuscript. All authors approved the final version of the manuscript.

## Acknowledgements

The authors wish to thank members of the Alexander lab including Jeffrey Fairley and Grace Morrison for their assistance with experiments. The authors wish to thank Michael Lopez and Anna Thalacker-Mercer for their critical evaluation of the manuscript prior to submission. The authors wish to acknowledge Kirk Habegger, Shelly Nason, Jessica Antipenko, and members of the UAB Diabetes Animal Physiology Core for assistance with the metabolic experiments. The authors wish to acknowledge Mary Ballestas and Reid Millican from the UAB Neuroscience NINDS Vector and Virus Core (Funded by NINDS P30 grant number NS047466). Research reported in this publication was supported by Eunice Kennedy Shriver National Institute of Child Health and Human Development, NIH, HHS of the National Institutes of Health under award number R01HD095897 awarded to M.S. Alexander. M.S. Alexander is supported by an NIH Office of Research Infrastructure Program (ORIP) U54 grant U54OD030167. A.S. was funded by a NIH NINDS T32 training grant number 5T32NS095775.

